# Strategies for robust, accurate, and generalizable benchmarking of drug discovery platforms

**DOI:** 10.1101/2024.12.10.627863

**Authors:** Melissa Van Norden, William Mangione, Zackary Falls, Ram Samudrala

## Abstract

Benchmarking is essential for the improvement and comparison of drug discovery platforms. We revised the protocols used to benchmark our Computational Analysis of Novel Drug Opportunities (CANDO) multiscale therapeutic discovery platform to bring them into strong alignment with best practices.CANDO ranked 7.4% and 12.1% of known drugs in the top 10 compounds for their respective diseases/indications using drug-indication mappings from the Comparative Toxicogenomics Database (CTD) and Therapeutic Targets Database (TTD), respectively. Better performance was weakly correlated (Spearman correlation coefficient *>*0.3) with the number of drugs associated with an indication and moderately correlated (coefficient *>*0.5) with intra-indication chemical similarity. There was also a moderate correlation between performance on our original and new benchmarking protocols. Higher performance was observed when using TTD instead of CTD when drug-indication associations appearing in both mappings were assessed. CANDO is available at https://github.com/ram-compbio/CANDO. The version used in this paper is available at http://compbio.buffalo.edu/data/mc_cando_benchmarking2. Supplementary data, drug–indication interaction matrices, and drug–indication mappings are available at http://compbio.buffalo.edu/data/mc_cando_benchmarking2.

## 2 Introduction

Drug discovery is difficult: according to a 2010 estimate, 24.3 early “target-to-hit” projects were completed per approved drug. [1] These preclinical projects were estimated to account for between 31% and 43% of total drug discovery expenditure. [1, 2] The result is a high and increasing price for novel drug development, with estimates ranging from $985 million to over $2 billion for one new drug to be successfully brought to market. [2–4] The creation of more effective computational drug discovery platforms promises to reduce the failure rate and increase the cost-effectiveness of drug discovery. [5, 6] Thousands of articles have been published on this topic, and multiple drugs discovered and/or optimized through computational methods are already in use. [7, 8] Modern drug discovery techniques range from traditional single-target molecular docking and retrospective clinical analysis to newer signature matching, network/pathway mapping, and deep learning pipelines. [9–11] The successes and failures of novel and repurposed therapeutics in combating the COVID-19 pandemic made more clear than ever that robust and effective drug discovery pipelines are essential for modern healthcare. [11–14] Still, systems for the assessment and comparison of these computational platforms need improvement and standardization [15].

For this study, we define a drug discovery platform as consisting of one or more pipelines, themselves comprising protocols (individual processes like predicting drug-target interactions), that come together to predict novel drug candidates for different diseases/indications. Benchmarking is the process of assessing the utility of such platforms, pipelines, and protocols. [16, 17] Quality benchmarking assists in (1) designing and refining computational pipelines; (2) estimating the likelihood of success in practical predictions; and (3) choosing the most suitable pipeline for a specific scenario. Only a few publications have explored guidelines for benchmarking novel drug discovery platforms. [15, 18] Other key publications include those that created datasets or databases that are used for benchmarking, such as Cdataset, PREDICT, and LRSSL.[19–21] Such static datasets are used instead of or alongside continuously updated databases like Drugbank, the Comparative Toxicogenomics Database (CTD), and Therapeutic Targets Database (TTD). [22–24] The limited availability of guidance on drug discovery benchmarking alongside an abundance of data sources for said benchmarking has resulted in the proliferation of numerous different benchmarking practices across different publications. [15, 18]

Most drug discovery benchmarking protocols start with a ground truth mapping of drugs to associated indications, though numerous “ground truths” are currently in use. [18] Data splitting is also frequently required. K-fold cross-validation is very commonly employed. [19–21, 25–69] Training/testing splits, leave-one-out protocols, or “temporal splits” (splitting based on approval dates) are also used occasionally. [44, 70–73] Results are then encapsulated in various metrics. [15] Area under the receiver-operating characteristic curve (AUROC) and precision-recall curve (AUPR) are commonly used. [19–21, 25–33, 35–71, 74–87] However, their relevance to drug discovery has been questioned. [15, 80, 88] Interpretable metrics like recall, precision, and accuracy above a threshold are also frequently reported. [20, 21, 30–33, 35–41, 70, 81, 82, 89–92] Case studies often appear alongside quantitative assessments to provide a tangible confirmation of predictive power. [21, 30–33, 35–54, 56–64, 70, 80, 82–87, 93–100] This wide variation in benchmarking can make determining best practices for assessing a given drug discovery protocol difficult, but ensuring a maximally informative and minimally biased assessment remains a necessity.

We developed the Computational Analysis of Novel Drug Opportunities (CANDO) platform based on the hypothesis that drugs with similar multitarget protein interaction profiles have similar biological effects. [15, 73, 101–116] CANDO calculates all-against-all interaction signature similarities to predict drug candidates, and it has been extensively validated. [15, 73, 110–115, 117–126] Previous efforts to benchmark CANDO have focused on assessing its ability to generate useful drug-drug similarity lists, which are used to generate novel drug predictions. [15, 73, 101, 116]

Our goal for this study is to bring the benchmarking protocols of our drug discovery platform into strong alignment with best practices. We thus updated our internal benchmarking protocol to evaluate the predictions generate by CANDO based on a consensus of the previously-assessed similarity lists. This revision allowed us to optimize parameters used in CANDO and examine the influence of certain features on its performance. Utilizing the updated protocols and parameters thus created will result in improved performance.

## 3 Methods

### 3.1 Drug discovery using CANDO

The CANDO multiscale drug discovery platform predicts novel compounds for diseases/indications based on their multitarget interaction signatures. It has been described extensively in other publications. [15, 73, 101–105, 110, 111, 116] CANDO consists of multiple pipelines designed for different prediction scenarios; some require human prioritization of drug targets, like our multitarget screening pipeline, and others require that the indication being predicted for is associated with at least one drug, like the primary pipeline of CANDO. We assessed this latter pipeline in this article. In the primary pipeline, the interaction signatures of every compound are compared to those of every other compound under the hypothesis that compounds with similar signatures exhibit similar behaviors. Each compound is thus associated with a sorted “similarity list” containing every other compound ranked by signature similarity. We calculated compound-compound signature similarity as the root mean squared distance between their proteomic interaction signatures (vectors of compound-protein scores) in this study using rapid, parallelizable algorithms from scikit-learn. [73, 127]

CANDO combines multiple similarity lists into novel drug predictions for an indication via a consensus protocol: (1) The similarity lists of drugs associated with the indication are examined. (2) The most similar compounds to each associated drug are ranked. (3) A consensus score is assigned to each compound based on the number of similar lists in which it appears above a certain rank (the similarity list cutoff). Average rank in those lists is also calculated. (4) Compounds are sorted by consensus score and average rank above the similarity list cutoff. The best ranked compounds in this consensus list are the top predictions for an indication. This pipeline is summarized in Figure 1.

**Figure 1:**
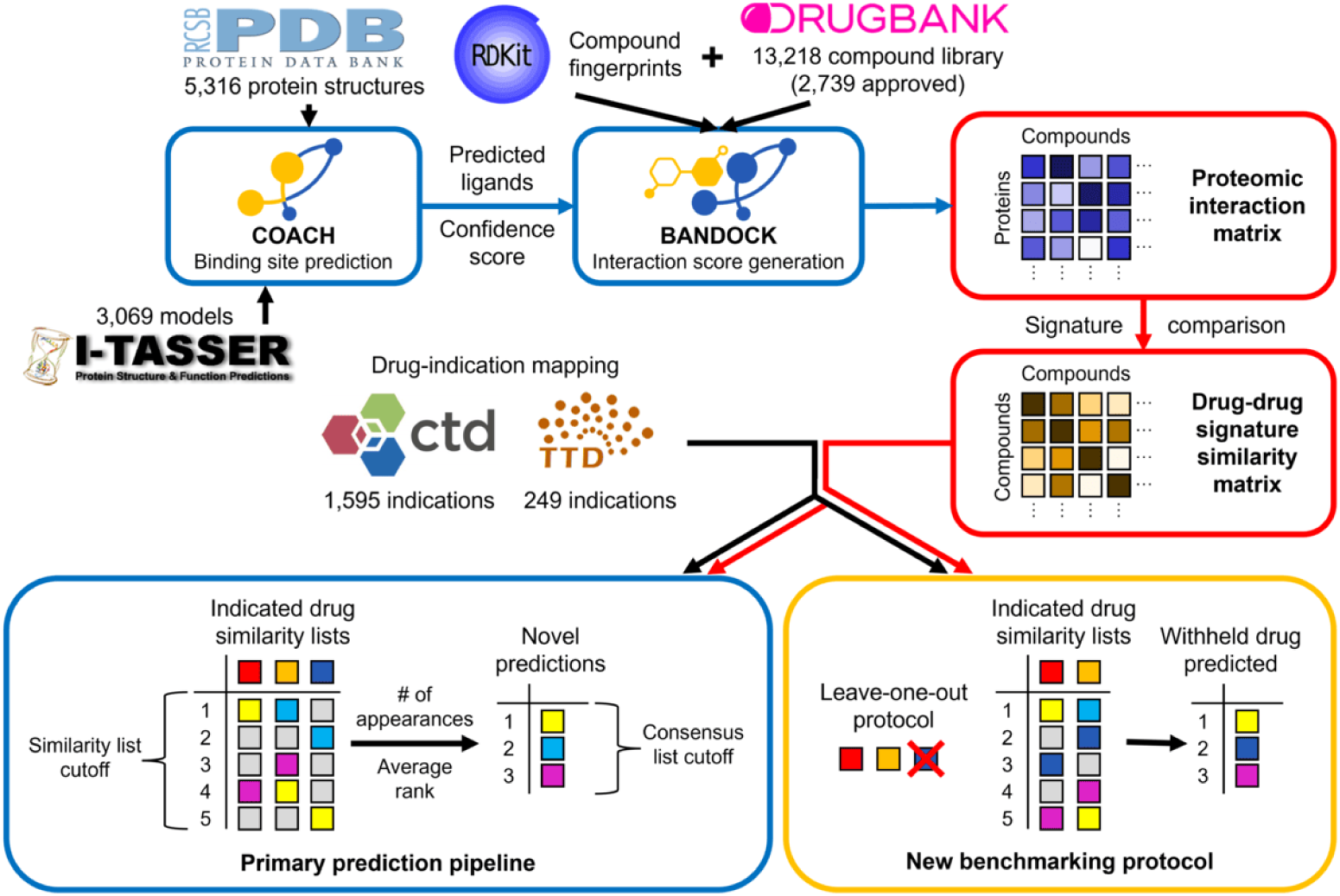
Flowchart of CANDO prediction and benchmarking pipelines. The primary prediction pipeline of CANDO is shown. Data sources are represented by their respective logos, protocols by blue boxes, key data structures by red boxes, and the benchmarking protocol by a yellow box. COACH is used to predict protein binding sites based off of experimental structures from the Protein Data Bank (PDB) and/or computational models created via I-TASSER. [128–131] Predicted ligands and confidence scores for each binding site are combined with compound fingerprints (from RDKit) to calculate protein interaction scores for every small molecule in the compound library (from DrugBank) using the bioanalytic docking (BANDOCK) protocol. [24, 132] These interaction scores are arranged into interaction signatures for every compound. Drug-drug signature similarity scores are calculated from these signatures. Drug-indication mappings are extracted from the CTD and/or TTD, and the most similar compounds to each drug associated with an indication are examined. Our original benchmarking protocol assessed the resulting similarity lists. Novel compound predictions are generated and ranked based on the number of times a compound appears in these lists above the similarity list cutoff; ties are broken based on average rank in these lists. In the example, the yellow compound is first because it appears the most times, and the cyan compound is second because its average rank is better than that of the magenta compound. The new benchmarking protocol uses the same scoring as the prediction pipeline, but each compound is left out in turn from its respective indication(s). The dark blue compound is left out of this example and then predicted with a rank of two based on the remaining indicated drugs. The original and new benchmarking protocols differ in what is assessed: the original focuses on the individual similarity lists, whereas the new evaluates the final consensus list.

### 3.2 Data extraction and generation

Proteomic interaction signatures were created based on the CANDO version 2.5 compound and human protein libraries. The protein library comprised 8,385 nonredundant human protein structures, including 5,316 experimentally determined structures extracted from the Protein Data Bank and 3,069 models generated using I-TASSER version 5.1. [102, 128–131]. Our bioanalytic docking (BANDOCK) protocol uses binding site data to calculate compound-protein interaction scores. We used the COACH pipeline to generate these data for our protein library. [133] COACH compared potential binding sites with solved bound protein structures to calculate binding site similarity scores and likely ligands. [112, 133] The chemical similarity between each compound in our library and the most similar predicted ligand was calculated using Extended Connectivity Fingerprints with a diameter of 4 (ECFP4) generated by RDKit. [112, 116, 132] Compound-protein interaction scores were then calculated in three ways: (1) as the chemical similarity score (the compound-only or C score), (2) as the product of the chemical similarity score and the binding site similarity score (the compound-and-protein or CxP score), or (3) as the product of the percentile chemical similarity score and the protein binding score (the percentile compound-and-protein or dCxP score). We compared all three interaction scoring types in our parameter optimization study (Methods: Optimizing parameters); the second score was used for our predictive power assessment.

Benchmarking requires known drug-indication mappings, which we obtained from two sources. We combined drug approval data from DrugBank and drug-indication associations from the CTD to make the “CTD mapping,” which is also available in version 2.5 of CANDO. [22, 24] The “TTD mapping” was created from approved drug-indication associations downloaded from the TTD; only compounds with existing interactions signatures were included. [23] In total, there were: 2,449 approved drugs across 2,257 indications with at least one associated drug and 22,771 associations in the CTD drug-indication mapping; 1,810 drugs across 535 indications and 1,977 associations in the TTD mapping; and 2,739 unique drugs altogether. Of these indications, 1,595 were associated with at least two drugs and thus could be benchmarked in CTD, and 249 were associated with at least two drugs in TTD.

### 3.3 Benchmarking CANDO

Our original benchmarking protocol examined the similarity lists of each indication-associated drug. [15, 73, 101–116] Indication accuracy (IA) was calculated as the percentage of these similarity lists in which at least one associated drug appeared above a cutoff. Indication accuracies were then averaged for every assessed indication to obtain average indication accuracy (AIA).

Our new benchmarking protocol uses the Python modules polars and sqlalchemy to rapidly evaluate the consensus scoring protocol and more accurately measure the performance of CANDO. [134, 135] To assess an indication, each associated drug is withheld from the indication in turn. Compounds are ranked as in the consensus scoring protocol (Methods: Drug discovery using CANDO). Two additional tiebreakers are used to order compounds that fall outside of the similarity list cutoff: (1) average rank across the full similarity lists of indication-associated drugs and (2) average similarity to associated drugs. The rank of the withheld drug in the final sorted list is determined and used to calculate multiple metrics.

Our new protocol calculates two primary metics. New indication accuracy (nIA) is calculated as the percentage of withheld drugs that are predicted at or above a defined rank cutoff in the consensus list. We chose rank cutoffs of 10, 25, and 100 for this study. nIA is averaged across all assessed indications to calculate new average indication accuracy (nAIA). Second, normalized discounted cumulative gain (NDCG) prioritizes early discovery of true positives. [15] We calculate discounted cumulative gain as follows:

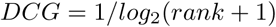

This is divided by the ideal discounted cumulative gain (here, equal to one) to obtain NDCG. Greater NDCGS are better. This metric will be referred to as new NDCG (nNDCG) when calculated by our new protocol. We determined nNDCG without a rank cutoff (overall) and at rank cutoffs of 10, 25, and 100 in this study.

### 3.4 Optimizing parameters

We optimized two CANDO parameters with regards to performance on nAIA and nNDCG. We randomly split the indications in our drug-indication mappings 30 to 70 to create independent mappings for parameter optimization and performance evaluation, limiting the risk of bias in our final assessment. All associations with the same indication were assigned to the same group, and only indications with at least two associated drugs were assessed. The CTD mapping was split into 5,714 drug-indication associations across 501 indications for parameter optimization and 13,226 associations across 1,094 indications for the final assessment. The smaller TTD mapping was split into 490 associations across 82 indications for parameter optimization and 1,160 associations across 167 indications for the final assessment.

The first parameter optimized was the similarity list cutoff used in our consensus scoring protocol (Methods: Drug discovery using CANDO). We quantified performance using nAIA and nNDCG on the CTD and TTD drug-indication mappings for every value of this parameter up to the number of approved drugs in the mapping (2,449 for CTD and 1,810 for TTD). The optimal parameter was the similarity list cutoff used when each metric reached its maximum. A random control was calculated for each metric and mapping at each optimal parameter. A hypergeometric distribution was used to calculate the control nAIA. For nNDCG, ten randomized drug-protein interaction matrices were generated and benchmarked per optimal value and mapping, and the nNDCGs were averaged.

The second parameter optimized was the compound-protein interaction scoring type. We benchmarked CANDO using proteomic interaction matrices generated using each of the three BANDOCK scoring types (Methods: Data extraction and generation) with similarity cutoffs ranging from 1 to 100. We compared the best performances of each scoring type using nAIA and nNDCG.

### 3.5 Evaluating features affecting performance

A final assessment was completed using the 70% of indications not used for parameter optimization. Similarity list cutoffs of six, ten, and thirteen and the compound-and-protein interaction score were used based on the parameter optimization results of both mappings (Methods: Optimizing parameters). We calculated nAIA and nNDCG at rank cutoffs of 10, 25, and 100, in addition to overall nNDCG, in this final assessment. We examined how three features correlated with performance, as quantified by nIA: the number of drugs associated with an indication, similarity list quality as quantified by our previous benchmarking metric (IA), and the chemical similarity of indication-associated drugs. Spearman correlation was used as our metrics do not follow a normal distribution, violating the assumptions of Pearson correlation. Correlation coefficients were calculated using the scipy package. [136] IA was generated by our original benchmarking protocol (Methods: Benchmarking CANDO). [73] Drug-drug chemical signature similarity was calculated using the Tanimoto coefficient on 2048-bit ECFP4 vectors, which encode the chemical features of a compound. ECFP4 fingerprints were generated by RDkit.[132, 137] Three similarity metrics were calculated for each indication: best similarity between any pair of associated drugs, average of the best similarities of each associated drug, and average of the average similarities of each associated drug.

### 3.6 Comparing drug-indication mappings

We compared benchmarking performance when using the mappings extracted from CTD and TTD. We combined the drugs from both mappings into a single library and benchmarked CANDO using this library and each full mapping. We manually matched each TTD indication to the most similar CTD indication. TTD indications were excluded if no appropriate CTD indication existed or if the most appropriate CTD indication was already matched to a more similar TTD indication. Performance on each mapping, quantified as nAIA, was compared for the matched indications. Finally, we compared the rankings of the drugs that were associated with the same indications in both CTD and TTD.

## 4 Results and discussion

In this study, we created two new benchmarking protocols to allow more consistent assessment of CANDO and computational drug discovery platforms in general. We present results obtained via these new protocols, including (1) the optimization of key parameters involved in our primary prediction pipeline; (2) an assessment of CANDO using these optimized parameters, including the correlations between performance on the new benchmarking protocol and the number of drugs associated with a disease/indication, performance on our original benchmarking protocol, and the drug-drug chemical signature similarity within an indication; and (3) a comparison of performance using two different drug-indication mappings.

### 4.1 Optimization of two key CANDO parameters

Our new internal benchmarking protocol allows us to directly assess the performance of the consensus scoring protocol CANDO uses to rank potential therapeutics (Methods: Benchmarking CANDO). This allowed us to optimize a key parameter of this protocol. CANDO uses a similarity list cutoff to determine how many similar compounds are considered per indication-associated drug (Methods: Drug discovery using the CANDO platform). Various similarity list cutoffs have been used in previous applications of CANDO. [108–110, 113–115]

We benchmarked the performance of CANDO at every possible similarity list cutoff on subsets of drug-indication mappings extracted from the CTD and TTD. Results were quantified using new average indication accuracy (nAIA) and new normalized discounted cumulative gain (nNDCG) at the top10, top25, and top100 rank cutoffs; nNDCG was also calculated without a rank cutoff. The results for similarity list cutoffs up to 1,810 are shown in Figure 2.

**Figure 2:**
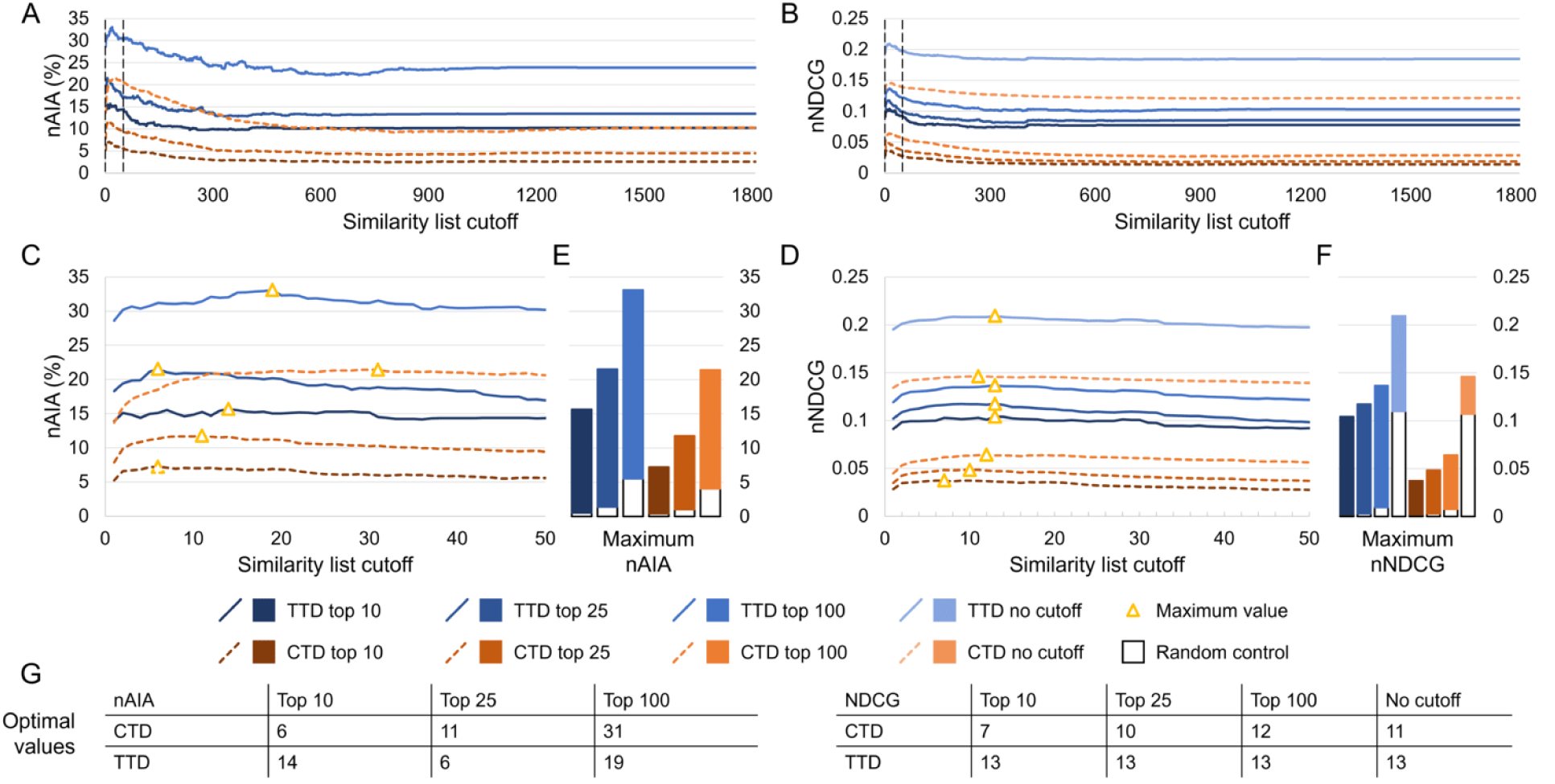
Effects of the similarity list cutoff on benchmarking performance. We used our new bench-marking protocol to optimize the similarity list cutoff, which represents the number of similar compounds the consensus protocol considers per associated drug when predicting a new compound for an indication. Assessments were completed on two drug-indication mappings extracted from the CTD and TTD. Results were summarized using nAIA and nNDCG metrics at multiple rank cutoffs; nNDCG was also calculated without a rank cutoff. Performance using nAIA (A) and nNDCG (B) is shown for similarity list cutoffs up to 1,810. Dotted black lines indicate the cutoffs of 1 and 50, between which all optimal values for this parameter fall. An expanded graph of only this range is shown for nAIA (C) and nNDCG (D). Bar charts (E–F) show the maximum values of each metric against random controls. Optimal values are marked with a yellow triangle and listed in the tables (G) at the bottom. The optimal parameter values for nAIA varied from 6 to 31 based on the cutoff and mapping used. The range was smaller for nNDCG, ranging from 7 to 13. The similarity list cutoff affected performance on multiple key metrics, and optimal performance was only achieved when less than 2% of compounds were considered.

Performance varied based on the similarity list cutoff used. The largest range was observed in the CTD mapping using nAIA top100, which ranged from 9.2% (similarity list cutoff of 805) to 21.4% (cutoff of 31). CANDO outperformed the random control across all metrics and similarity list cutoffs. Optimal parameter values ranged from 6 (nAIA top10 using CTD and nAIA top25 using TTD) to 31 (nAIA top100 using CTD). Performance was better on all metrics and cutoffs when using the TTD mapping instead of the CTD mapping.

We also assessed the effect of the protocol used to calculate drug-protein interaction scores. Our BANDOCK interaction scoring protocol computes three types of interaction scores: compound-only, compound-and-protein, and percentile compound-and-protein (Methods: Data extraction and generation). We optimized the similarity list cutoff using nAIA and nNDCG with all three interaction scoring types; the best performances of each scoring type on each metric were compared (Supplementary Table 1). The compound-and-protein score showed the best performance on most metrics when using the CTD mapping, with the percentile compound-and-protein score performing best on the remaining metrics. On the other hand, the compound-only score performed best on the majority of metrics when using the TTD mapping, with the compound-and-protein score performing best on one. We thus determined that the compound-and-protein score was the best scoring type for our uses as it was often the best performing and never the worst performing type.

### 4.2 Assessment of predictive power

We conducted a second assessment to determine the predictive power of CANDO using our optimized parameters. We conducted three assessments on the drug-indication associations not used for optimization using the compound-and-protein interaction score and similarity list cutoffs of six, ten, and thirteen, chosen based on our optimization trials. The results are shown in Figure 3A–B.

**Figure 3:**
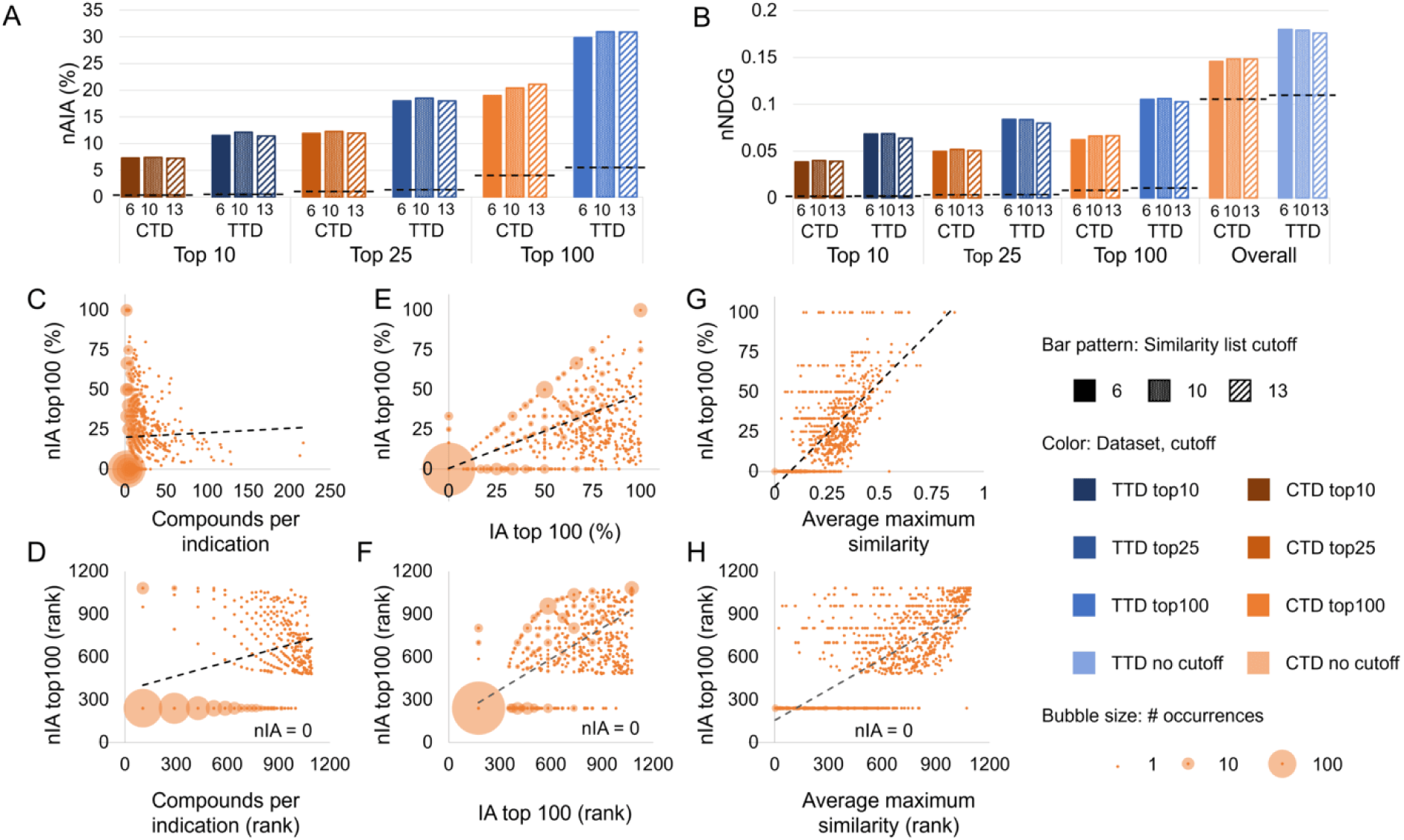
Assessment of predictive power. CANDO was assessed using the protocols and parameters obtained through our optimization. nAIA (A) and nNDCG (B) metrics are shown at multiple cutoffs for the two drug-indication mappings, CTD and TTD. The random control is shown as a dotted line on each group of bars with the same mapping and cutoff. CANDO outperformed the control on all assessments, and performance was best when using the TTD mapping. Performance on this assessment was correlated with multiple features: the number of compounds in an indication (C–D); our original indication accuracy (IA) metric, which measures similarity list quality (E–F); and drug-drug chemical signature similarity within each indication, measured as the average Tanimoto coefficient between the chemical fingerprint of each drug and that of its most similar other associated drug (G–H). The upper subfigures (C, E, and G) plot each feature against nIA top100 in CTD, whereas the lower plots (D, F, and H) show the relationship between the same two features when their values are ranked; these ranks were used to calculate Spearman correlation coefficients. The size of the bubble surrounding each dot represents the number of indications plotted there. Trendlines are shown as dotted black lines. Positive correlations of varying strength were observed in all cases. Knowledge of the features influencing benchmarking can enable more accurate assessment of expected predictive performance. Based on these results, we can expect CANDO to perform best when predicting compounds for indications with large numbers of associated drugs and high chemical signature similarity.

CANDO outperformed random controls using both drug-indication mappings and all metrics. nAIA results showed that CANDO recovered 7.3% to 7.4% of approved drugs within the top 10 compounds using the CTD mapping and 11.4% to 12.1% using the TTD mapping (out of 2,449 compounds in CTD and 1,810 in TTD). This rose to 19.0% to 21.1% using CTD and 29.9% to 31.0% using TTD at the top 100 cutoff. nNDCG top10 ranged from 0.038 to 0.040 using CTD and 0.064 to 0.068 using TTD, more than an order of magnitude greater than the corresponding random control values. Data for all metrics and similarity list cutoffs are available in Supplementary Table 2.

#### 4.2.1 Influence of the number of associated drugs

We investigated three features that could influence the performance of CANDO to understand what makes it perform better on some indications than others. First, we considered the influence of the number of drugs associated with an indication (indication size) on performance.

Greater data availability generally improves the performance of computational models, so we anticipated a positive correlation between nIA and indication size. There was indeed a weak positive correlation, with Spearman correlation coefficients ranging from 0.324 to 0.352 using the CTD mapping and from 0.337 to 0.505 using the TTD mapping. The correlation between nIA at the top100 cutoff and indication size using the CTD mapping is illustrated in Figure 3C–D. Correlations between nIA at all cutoffs are shown in Supplementary Figure 1.

We wondered if this apparent positive correlation could result from the large number of indications with few associated drugs that had an nIA of zero. We thus re-calculated the correlation coefficient using only indications with five or more associated drugs (Supplementary Table 3A). The strength of the correlation between nIA and indication size weakened in this assessment, becoming negligible at 0.007 to 0.071 using CTD and shrinking to 0.057 to 0.252 using TTD. This suggests that the benefit of increased indication size is limited beyond five drugs. This could result from indications with more associated drugs having more associated mechanisms of action and disease subtypes (e.g., HER2-positive or triple negative breast cancer), decreasing the similarity of the associated drugs.

#### 4.2.2 Influence of similarity list quality

We also considered the primary metric of our previous internal benchmarking protocol: indication accuracy (IA; Methods: Benchmarking CANDO). IA directly measures the quality of the drug-drug interaction signature similarity ranks calculated by CANDO. IA is more lenient and thus tends to be higher than nIA: IA checks whether at least one other associated drug appears above a certain cutoff rather than the percentage of associated drugs recalled.

We found a strong relationship between nIA and IA at the top10, 25, and 100 cutoffs for each indication, with Spearman correlation coefficients ranging from 0.741 to 0.807 using the CTD mapping and 0.859 to 0.905 using the TTD mapping. No relationship was observed between the cutoff considered and the strength of the correlation. The correlation between nIA and IA at the top100 cutoff using the CTD mapping is illustrated in Figure 3C–D, and the remaining correlations are illustrated in Supplementary Figure 2. The correspondence between IA and nIA suggests that our previous benchmarking results did have relevance to actual performance, as has also been demonstrated by extensive prospective validation. [15, 110–115, 117–126] This also, unsurprisingly, suggests that high-quality similarity lists result in high-quality consensus predictions.

#### 4.2.3 Influence of chemical signature similarity

CANDO typically uses proteomic interaction signature similarity between indication-associated drugs and other compounds to predict whether those compounds would be effective for that indication. Compounds that are alike in chemical structure tend to have similar protein interactions and, therefore, similar interaction signatures. [116] Note that the use of only approved drugs confounds this assessment due to the relatively high presence of "me-too" drugs compared to small molecules as a whole.

We assessed the extent to which chemical similarity, calculated as the Tanimoto coefficient between the chemical fingerprints of two drugs, influences the performance of CANDO. [137] We quantified chemical similarity within an indication through three metrics: maximum similarity between any pair of associated drugs; average similarity across all pairs of associated drugs; and the average of the maximum similarities of each associated drug. The correlation was strongest using average maximum similarity, for which coefficients ranged from 0.635 to 0.750 using CTD and 0.697 to 0.744 using TTD (Supplementary Table 3C). The correlation between nIA top100 and average maximum similarity using the CTD mapping is illustrated in Figure 3G–H, and correlation with nIA top10, top25, and top100 using both mappings are shown in Supplementary Figure 3. Though the correlation was moderate-to-strong, the fact that the correlation was greatest when using average maximum similarity suggests that CANDO performs best when each drug has at least one chemically similar partner, but the indication does not need to be totally chemically homogeneous. This is supported by our previous work, which demonstrated that CANDO performs better when using the protein interaction signature than chemical similarity (aka molecular fingerprints), showing that the platform is not restricted to predicting compounds based on chemical similarity alone. [102, 116] CANDO has also demonstrated its ability to design and/or select compounds with novel chemical spaces when the appropriate inputs are used. [107]

### 4.3 Comparison of drug-indication mappings

CANDO consistently performed better when using the drug-indication mapping created from the TTD relative to the one from the CTD. There are a few reasons why this could occur. First, CTD and TTD contain different indications. If the indications in TTD are easier to predict drugs for than those in CTD, this could result in better benchmarking performance when using TTD. Second, CTD has more approved drugs in total, which means that each assessed drug has to outcompete more total compounds during benchmarking, decreasing performance. Finally, the TTD mapping has a higher standard of evidence for inclusion (FDA approval) than CTD (therapeutic association in the literature). This could lead to higher data quality and, thus, better performance.

We examined these drug-indication mappings head-to-head to determine whether using TTD actually improved performance. We benchmarked CANDO using both full mappings and the full library of drugs that are approved in either mapping. CANDO still performed better when using the TTD mapping, with a top10 nAIA of 6.8% using CTD compared to 11.3% using TTD. However, this difference decreased when only the 191 indications that appeared in both mappings were assessed. The top10 nAIA using CTD with matched indications was 6.5% compared to 9.3% using TTD. The differences in nIA top10 and top100 on each matched indication are shown in Figure 4A–B. Note that the CTD mapping performed better on more indications than the TTD, but the TTD generally outperformed by a greater magnitude.

**Figure 4:**
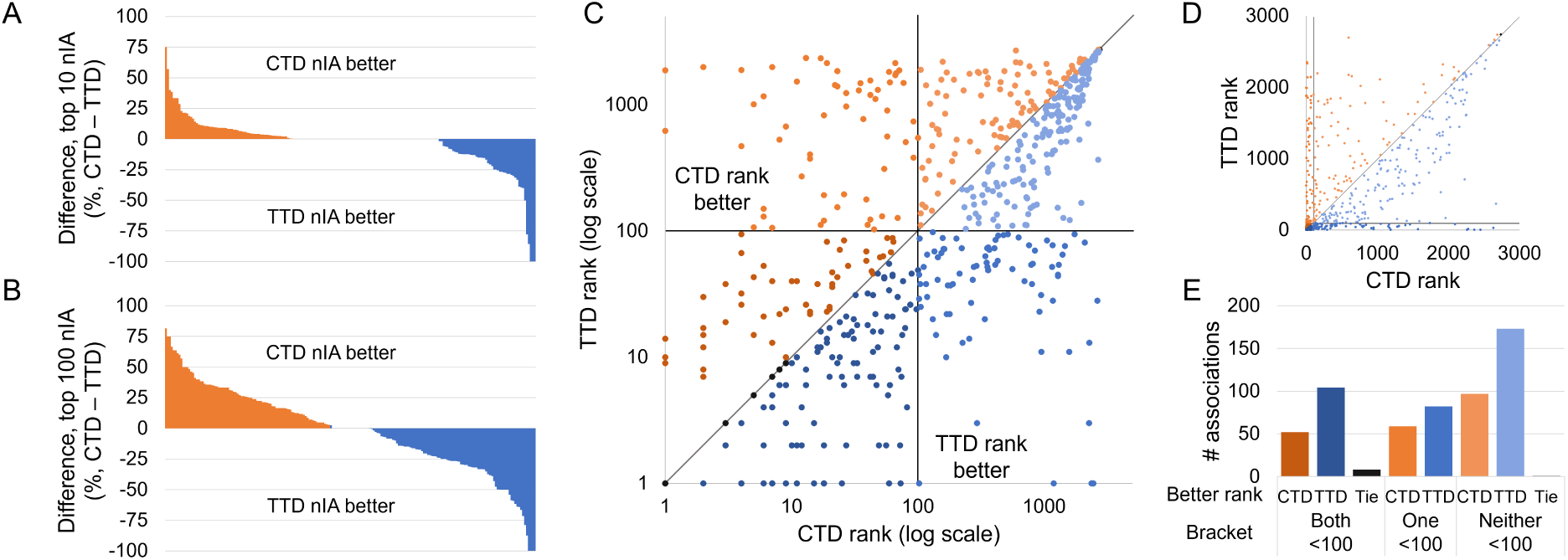
Influence of drug-indication mapping on performance. CANDO was benchmarked using drug-indication mappings extracted from two databases, CTD and TTD. The differences in nIA at the top10 (A) and top100 (B) cutoffs for each indication that appeared in both mappings are shown. Performance was better using CTD for more indications, but TTD outperformed by more when it was superior. This lead to a higher overall nAIA when using TTD. We also compared the ranks of 576 drug-indication associations that appeared in both mappings. The ranks of those drugs when predicted for their indications in each mapping are plotted in log scale (C) and arithmetic scale (D). Black lines indicate the 100*^th^* rank, beyond which predictions are less likely to be useful for drug discovery, and a grey line represents equivalent ranks between the mappings. The number of associations for which each mapping performed better is shown (E); these counts are separated by whether both, only one, or neither mappings ranked the drug within the top100 cutoff. A drug was more likely to rank well for its indication when using the TTD drug-indication mapping, but more individual indications performed better at the most stringent top10 cutoff when using the CTD mapping.

This trend was reversed when we grouped matched indications by the top-level MeSH headings they were associated with; TTD performed better on top10, top25, and top100 nAIA on 14, 13, and 15 out of 24 top-level MeSH headings, as compared to 7, 10, and 8 for CTD. Among those top-level groups containing at least 10 indications, TTD outperformed CTD by the most on “Endocrine System Diseases” (12 indications, top100 nAIA of 26.9% using CTD versus 46.3% using TTD), and CTD outperformed TTD by the most on “Infections” (32 indications, top100 nAIA of 37.6% using CTD versus 23.5% using TTD). The full results for this analysis are available in Supplementary Table 4.

The top10 nAIA was 21.5% greater when the full TTD mapping was used compared to only those indications matched with CTD indications. Meanwhile, there was only a 4.6% increase when the full CTD mapping was used over the matched indications. This indicates that the apparent better performance when using TTD may in part be due to its inclusion of “easier” indications that CANDO performs better on. Indications that were exclusively benchmarked using TTD with with high nIAs include anesthesia (ICD-11 9a78.6; 33 drugs, 15.1% top10 nIA), contraception (ICD-11 qa21; 10 drugs, 30% top10 nIA) and virus infection (ICD-11 1a24-1d9z; 8 drugs, 50% top10 nIA). However, this cannot be the only factor, as TTD still performed better when only matched indications were considered.

We next examined the 576 drug-indication associations that appeared in both CTD and TTD. The rankings our benchmarking protocol assigned to these drugs when using each mapping are plotted in Figure 4C–E. Of these drugs, 208 had better ranks using CTD, 359 had better using TTD, and 9 had the same ranks either way. Drugs were also more frequently ranked in the top10, 25, and 100 cutoffs when using TTD. CTD generally outperformed TTD by a greater magnitude when it was better, but, on average, matched drugs were predicted 23.5 ranks higher when using TTD.

We analyzed some specific drug-indication associations to further illustrate this difference. We first considered enzalutamide, an androgen receptor inhibitor approved for use in patients with prostate cancer. [138] CANDO ranked enzalutamide 44th for this indication using CTD, but its rank improved to 7th using TTD. Enzalutamide had the same consensus score across both mappings because it was among the top ten most similar compounds to the same drugs in both: bicalutamide, flutamide, and nilutamide. TTD included only 13 drugs in its prostate cancer indication, whereas CTD included 88 drugs. The greater number of associated drugs in the latter indication mapping thus led to increased consensus scores in general, making the score of enzalutamide less remarkable. The CTD prostate cancer indication included drugs like methotrexate, gemcitabine, and thalidomide that have shown efficacy for other cancers, but which are not approved for use in prostate cancer specifically. [139–145] CTD also contains a related indication: “Prostatic neoplasms, castration-resistant.” Enzalutamide, which is approved for use against such neoplasms, ranked second for this indication during benchmarking based on its similarity with bicalutamide and apalutamide. Therefore, the greater data availability of data in CTD can also be beneficial due to the existence of a greater quantity of more specific indications that may not be available in TTD.

Apalutamide is another drug approved for use in castration-resistant or metastatic castration-sensitive prostate cancer. [146] It was associated with castration-resistant prostate cancer in CTD, but neither mapping associated apalutamide with the general prostate cancer indication. We used the primary prediction pipeline of CANDO to fill this gap: it ranked apalutamide 81st for prostate cancer in CTD and 14th in TTD, providing additional evidence of the superiority of TTD even on associations not included in the original data. However, this also illustrates the limitations of the mappings used. No data source is perfect; if a perfect drug-indication mapping existed, drug discovery pipelines would be unnecessary. Still, such avoidable errors impact performance: enzalutamide would have ranked higher in both mappings had apalutamide been included. In addition, errors decrease the reliability of benchmarking, as platforms are judged on the ability to predict fictitious drug-indication associations while real relationships are left unassessed. Updates and quality control of drug-indication data is thus necessary to ensure our platforms are built on reliable data and our assessments remain as accurate as possible.

Internal benchmarking has shown that CANDO consistently outperforms random chance, particularly when using a drug-indication mapping consisting of approved associations only. CANDO was able to predict up to 31% of existing drugs in the top 100 potential therapeutics for their indications when using TTD. We were also able to optimize multiple parameters and elucidate the factors associated with better performance using our new benchmarking protocol. However, our current work is limited in that it is an internal bench-marking method, and it only allows limited comparison with other drug discovery platforms. Future work will therefore focus on comparable, head-to-head drug discovery benchmarking. Such in-depth and accurate benchmarking helps improve drug discovery platforms, ensures the reliability of published platforms, and maintains the quality of the field as a whole.

## Acknowledgments

The authors would like to acknowledge the Center for Computational Research of the University at Buffalo for their computational resources and support. We would also like to thank the members of the Samudrala Computational Biology Group.

## 6 Data and code availability

CANDO is publicly available through Github at https://github.com/ram-compbio/CANDO. Supplementary data, drug–indication interaction matrices, and drug–indication mappings are available at http://compbio.buffalo.edu/data/mc_cando_benchmarking2.

