## Supplementary Figures for "Strategies for robust, accurate, and generalizable benchmarking of drug discovery platforms"

### Supplementary figure 1 – Indication size

CTD, excluding other indicated compounds from rankings

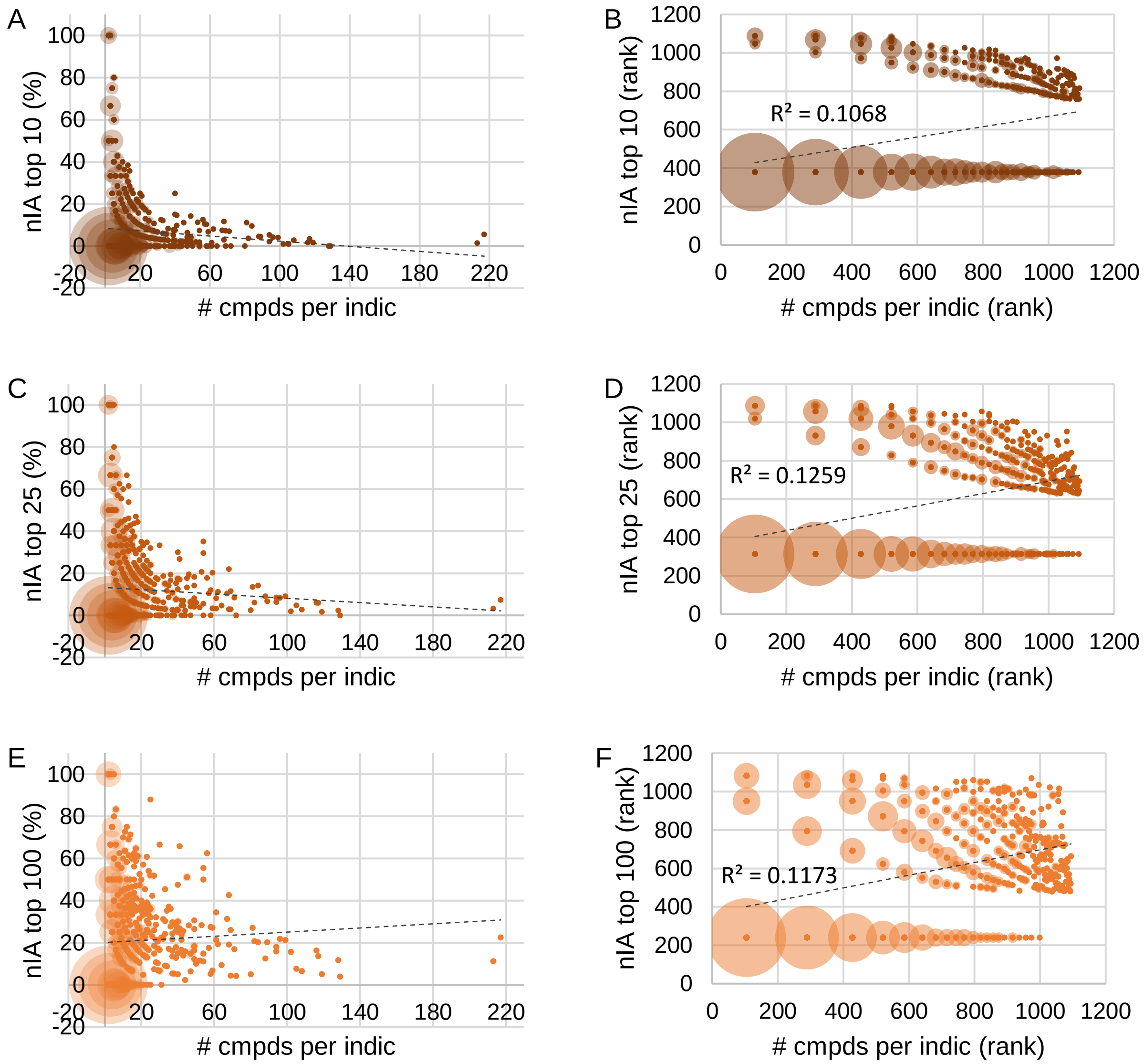


TTD, excluding other indicated compounds from rankings

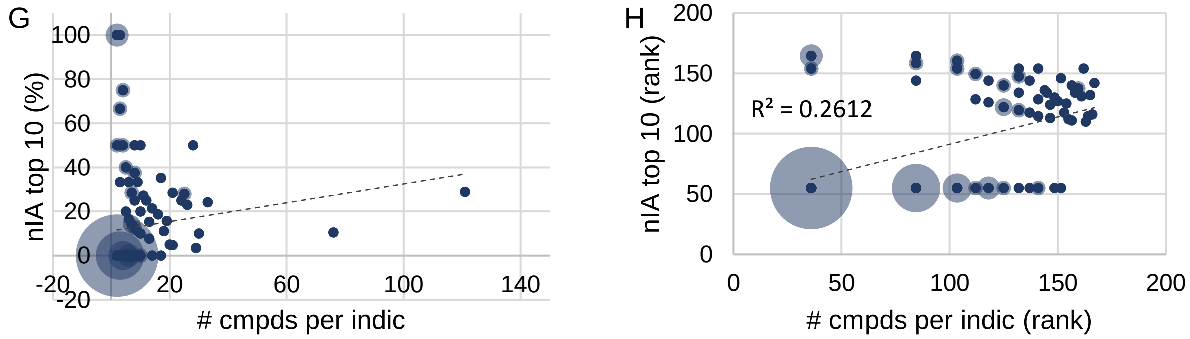

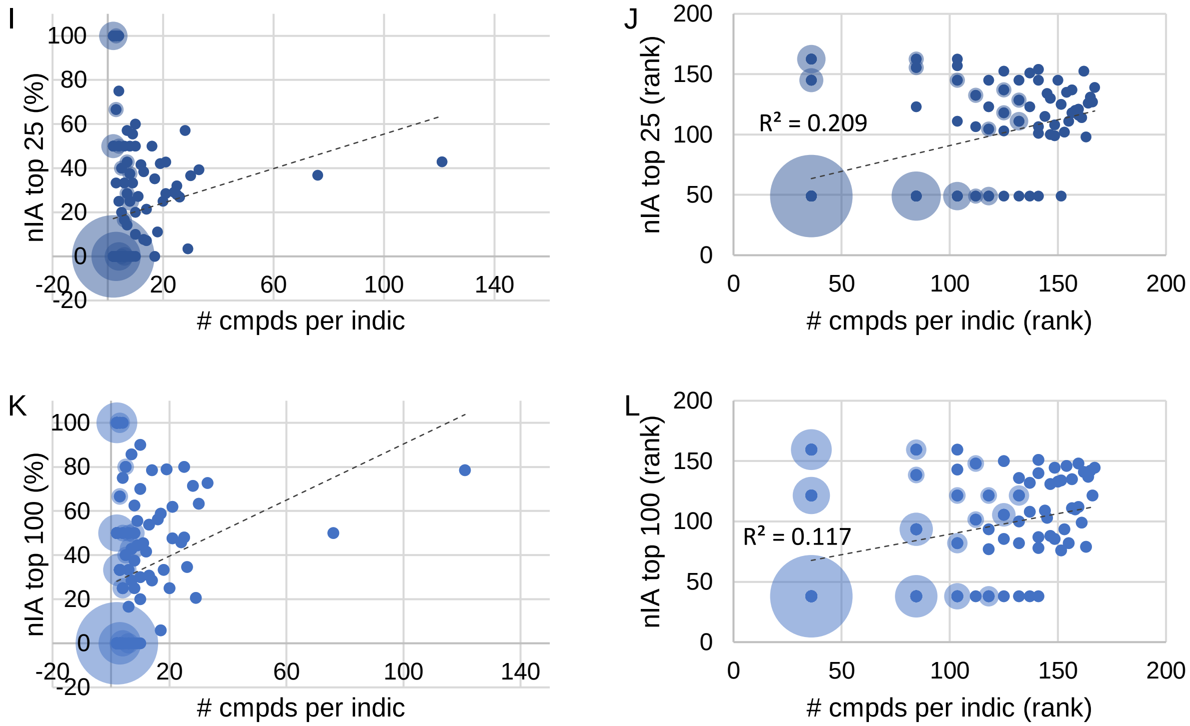
I
nIA top 25 (%)
### cmpds per indic
J
nIA top 25 (rank)
### cmpds per indic (rank)
R² = 0.209
K
nIA top 100 (%)
### cmpds per indic
L
nIA top 100 (rank)
### cmpds per indic (rank)
R² = 0.117

### Supplementary figure 2 – nIA versus IA

CTD

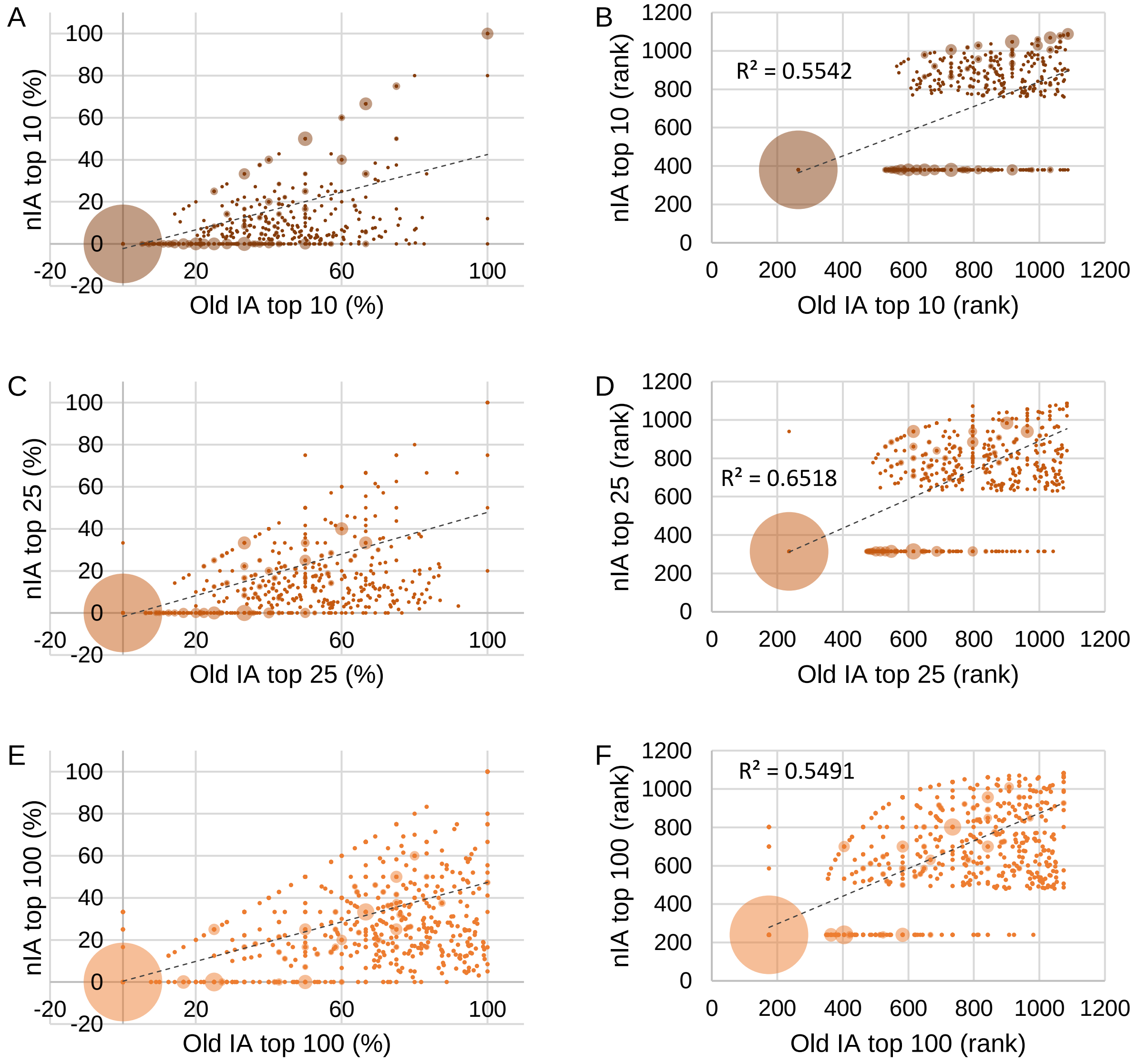


TTD

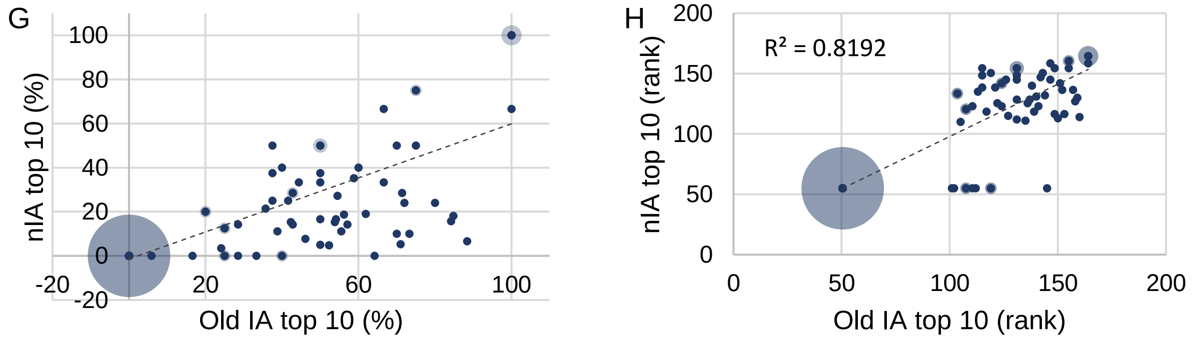

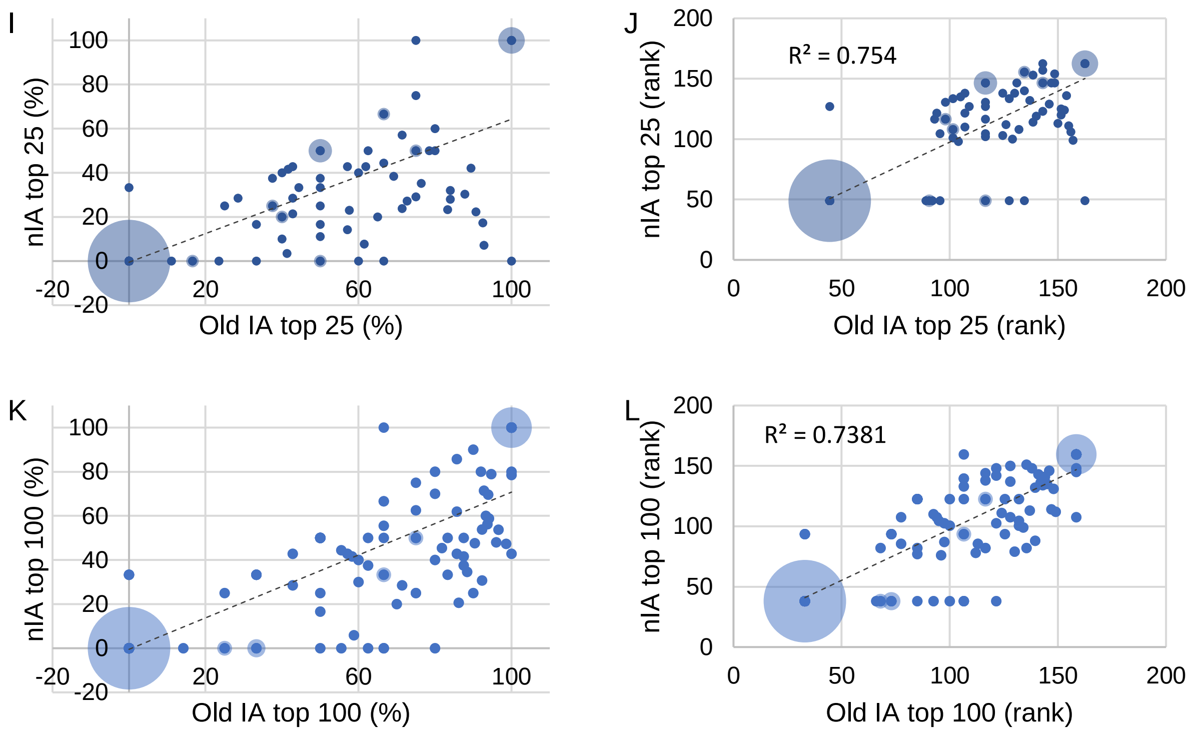
I
nIA top 25 (%)
Old IA top 25 (%)
J
R² = 0.754
nIA top 25 (rank)
Old IA top 25 (rank)
K
nIA top 100 (%)
Old IA top 100 (%)
L
R² = 0.7381
nIA top 100 (rank)
Old IA top 100 (rank)

### Supplementary figure 3 – Compound similarity

CTD

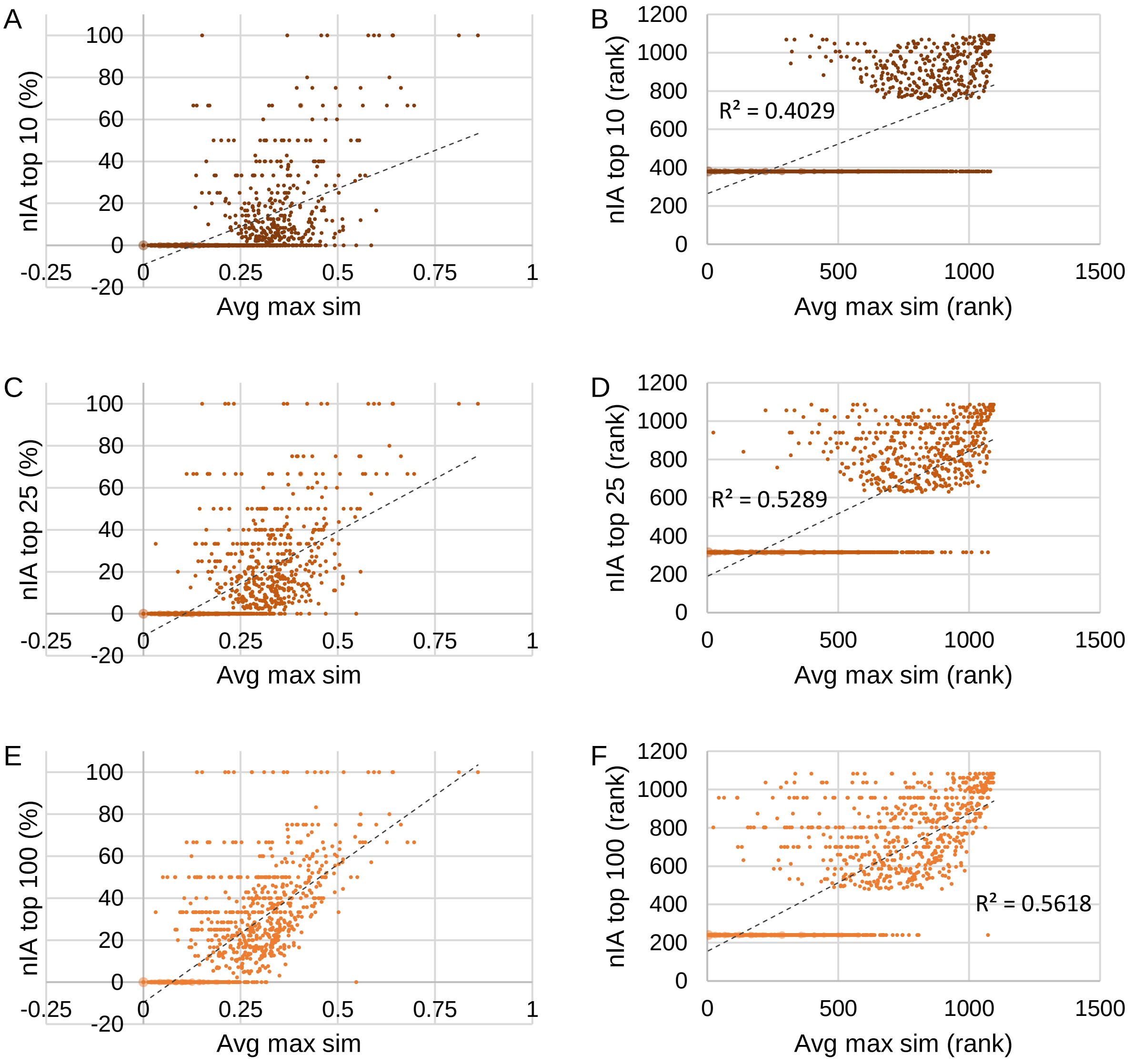


TTD

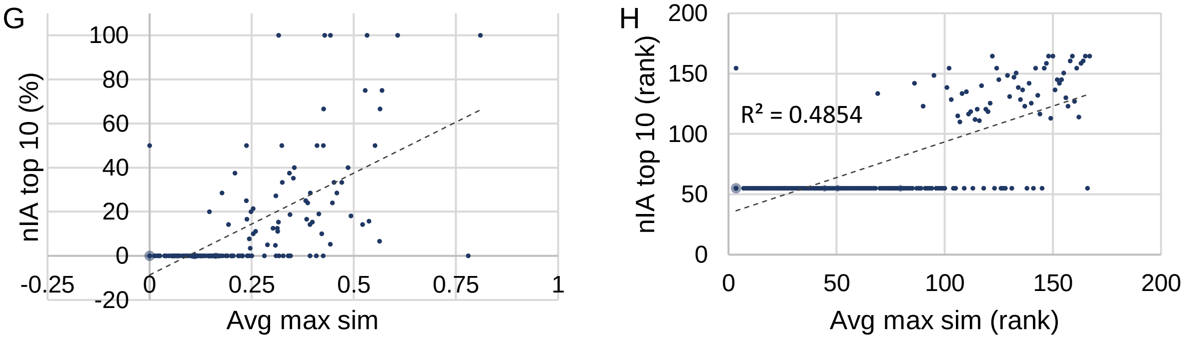

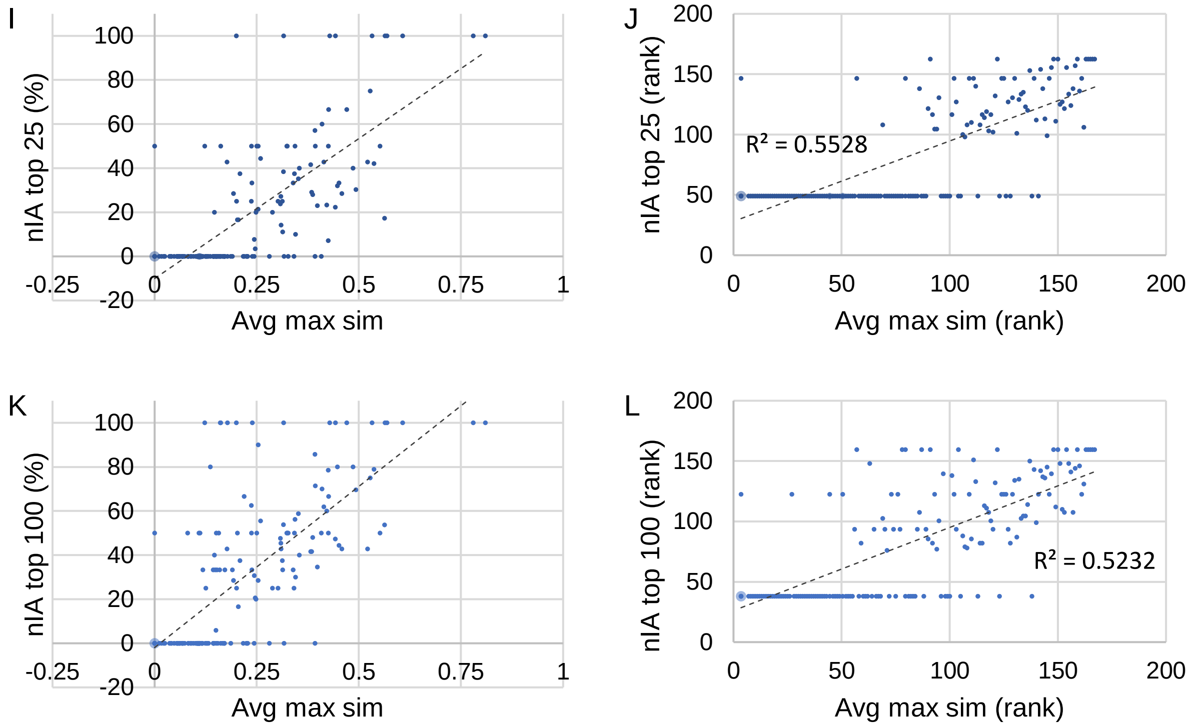

I
nIA top 25 (%)
Avg max sim
J
nIA top 25 (rank)
Avg max sim (rank)
R² = 0.5528
K
nIA top 100 (%)
Avg max sim
L
nIA top 100 (rank)
Avg max sim (rank)
R² = 0.5232
